# Multicenter reliability of electric field simulations: Evidence from a traveling-subjects study

**DOI:** 10.64898/2026.06.08.730792

**Authors:** Dayana Hayek, Daria Antonenko, Ulrike Grittner, Axel Thielscher

**Affiliations:** Department of Neurology, University Medicine Greifswald, Greifswald, Germany; Berlin Institute of Health, Charité University Medicine Berlin, Berlin, Germany; Institute of Biometry and Clinical Epidemiology, Charité University Medicine Berlin, Corporate, Member of Freie Universität Berlin, Humboldt-University Berlin and Berlin Institute of Health, Berlin, Germany; Department of Health Technology, Technical University of Denmark, Kongens Lyngby, Denmark; Danish Research Centre for Magnetic Resonance, Department of Radiology and Nuclear, Medicine, Copenhagen University Hospital Amager and Hvidovre, Hvidovre, Denmark

**Author notes:** Shared-first authorship (Dayana Hayek, Daria Antonenko).

## Abstract

Multicenter neuroimaging studies face challenges from scanner-related measurement biases that can obscure biological effects. Traveling-subject (TS) designs allow direct quantification of such biases by scanning the same participants across multiple sites. While reliability of structural, functional, and diffusion MRI has been established, the reliability of electric field simulations for transcranial brain stimulation has not been examined. We assessed inter-scanner and scan-rescan reliability of simulated electric field magnitudes in 10 participants scanned twice on each of five 3T Siemens scanners, using SimNIBS v4.1 to simulate focal tDCS across seven cortical and cerebellar targets. ICC values indicated good to excellent inter-scanner (0.86-0.97) and scan-rescan (0.94-0.98) reliability, with between-scanner variance not exceeding measurement error. Intra-individual segmentation variability substantially explained within-person differences in field magnitudes, whereas image quality did not predict segmentation variability. These results demonstrate that individualized electric field simulations are robust to scanner-related variation, supporting their use in multicenter brain stimulation research.

## Manuscript

Multicenter neuroimaging studies face the challenge that scanner hardware, pulse sequences and reconstruction methods differ substantially across sites. Such differences introduce measurement bias that can obscure or mimic biological effects. Traveling-subject (TS) designs provide a direct empirical way to quantify these biases: the same participants undergo repeated magnetic resonance imaging (MRI) sessions across multiple scanners, allowing a decomposition of variance into scanner site and subject components [1]. Good to excellent reliability has been shown for functional, structural, and diffusion tensor imaging [2]. Reliability of electric field simulations is of particular interest to multicenter trials on brain stimulation, but has not yet been delineated.

Here, we used a TS study design to assess how well imaging sequences and settings could be harmonized across five 3T Siemens MR scanners to enable reliable comparisons of simulated electric fields across sites. We established structural T1- and T2-weighted MRI protocols according to prior recommendations for obtaining stable whole-head segmentations [3, 4]. Data from 10 participants (5 females, mean age in years ± SD: 33 ± 8) was acquired, each undergoing two consecutive runs (scan-rescan) on five scanners in Germany (Berlin, Dortmund, Essen, Greifswald, Leipzig), totaling up to 100 measurements. Head segmentation and electric field estimation was done using SimNIBS v4.1 [5]. Selection of electrode montages and target regions was based on our ongoing study [6] and included visual-spatial (right occipitotemporal cortex, rOTC, left posterior parietal cortex, lPPC), language (left inferior frontal gyrus, lIFG), motor (left primary motor cortex, lM1, right cerebellum, rCB), and executive (left and right dorsolateral prefrontal cortices, lDLPFC, rDLPFC) regions. We simulated focal transcranial direct current stimulation using a 3×1 set-up and extracted field magnitudes from target regions. Additionally, quality metrics derived from MRIQC [7] and tissue volumes below the electrodes were extracted and compared. Inter-scanner and scan-rescan (test-retest) reliabilities were assessed with intra-class correlations coefficients (ICC) and variance components (Var; **Supplementary Material**).

Electric field magnitudes for 2 mA focal tDCS ranged between 0.08 and 0.45 V/m across targets, with individual target means ranging from 0.17 to 0.24 V/m (**Figure 1**). ICC values ranged from 0.86 to 0.97 (median: 0.94) for inter-scanner reliability and 0.94 to 0.98 (median: 0.97) for scan-rescan reliability, demonstrating good to excellent reliability. Variance component analysis revealed between-scanner variance of 0.8% to 7.1% (mean/SD: 2.1/2.3%) and scan-rescan variance (within-scanner variability) of 1.2% to 5.5% (mean/SD: 3.3/1.5%). Statistical comparisons showed that site-related variance components did not differ significantly from measurement error variance across all targets (all p’s > 0.45), indicating that scanner differences were not larger than measurement uncertainty within the TS design.

**Figure 1.**
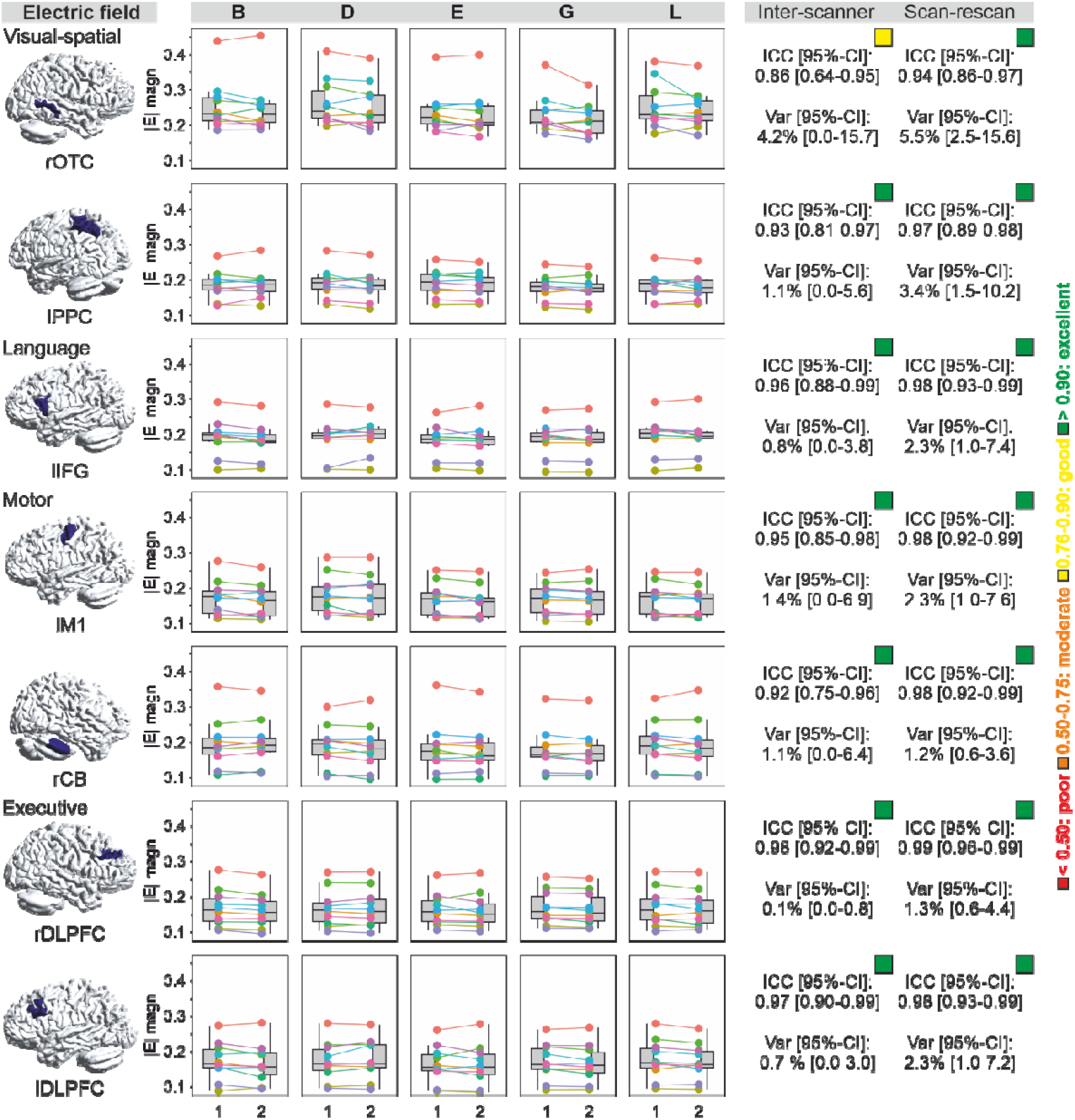
Electric field magnitudes for focal 2-mA tDCS set-ups, extracted from regions-of-interest, depicted in the left column (selected based on our multicenter study memoslap.de). The figures show two consecutive scans/runs for each of the 10 subjects (each in a different color) at five scanner sites (B, Berlin; D, Dortmund; E, Essen; G, Greifswald; L, Leipzig). Right columns show intra-class correlation coefficients (ICC) and variance components (Var) between sites (inter-scanner) and runs (scan-rescan). Colored squares above the ICC values additionally illustrate whether reliability was poor (ICC < 0.50, red), moderate (0.50-0.75, orange), good (0.76-0.90, yellow), or excellent (> 0.90, green). rOTC, right occipitotemporal cortex. lPPC, left posterior parietal cortex. lIFG, left inferior frontal cortex. lM1, left motor cortex. rCB, right cerebellum. r/lDLPFC, right/left dorsolateral prefrontal cortex.

Assuming that the observed differences, although small, derived from differences in structural segmentation, we also compared regional tissue volumes underneath the electrodes across scanners and scan sessions (i.e., volumes of scalp, bone, cerebrospinal fluid, and grey matter, **Supplementary Figures S1-S4**). ICC values demonstrated moderate to excellent reliability. Scalp volumes demonstrated inter-scanner ICC values from 0.84 to 0.96 (median: 0.93) and scan-rescan ICC from 0.94 to 0.98 (median: 0.97). Bone volumes exhibited inter-scanner ICC values from 0.85 to 0.96 (median: 0.93) and scan-rescan ICC from 0.93 to 0.98 (median: 0.96). CSF showed inter-scanner ICC values from 0.82 to 0.96 (median: 0.92) and scan-rescan ICC from 0.93 to 0.98 (median: 0.96). For GM, inter-scanner ICC values ranged from 0.73 to 0.94 (median: 0.87) and scan-rescan ICC values from 0.88 to 0.97 (median: 0.94). Variance components showed 0.9-10.2% between-scanner variance and 0.6-4.0% scan-rescan variance across tissue types.

In order to evaluate data quality, we compared signal-to-noise ratios (SNR), contrast-to-noise ratios (CNR), and entropy focus criterion (EFC) of T1 and T2 images (**Supplementary Figure S5**). ICC values demonstrated good to excellent reliability for most metrics: T1-weighted images (inter-scanner ICC: 0.74-0.92, scan-rescan ICC: 0.94-0.99) and T2-weighted images (inter-scanner ICC: 0.75-0.94 for SNR and CNR, scan-rescan ICC: 0.94-0.97). Variance components showed 0-3.1% inter-scanner variance and 0.9-3.1% scan-rescan variance. However, T2 EFC showed poor inter-scanner reliability (ICC = 0.26) with 44.0% between-scanner variance, indicating substantial differences in T2 image quality across sites despite protocol harmonization efforts due to different scanner software versions and sequence implementations (further details in **Supplementary Material**).

Total inter-scanner and scan-rescan variance within individuals in simulated electric field magnitudes was substantially explained by intra-individual differences in tissue segmentation (partial R^2^ values ranged between 0.44 to 0.71 across projects, **Supplementary Table S1**). Fixed effects revealed different contribution of intra-individual tissue differences, with scalp and CSF explaining the largest part of variance across all projects. Segmentation variability was, however, not explained by differenced in structural image quality (partial R^2^ values ranged between 0.001 to 0.18, **Supplementary Table S2**). These findings demonstrated that inter-scanner and scan-rescan differences in electric field magnitudes are primarily driven by segmentation variability, but that this variability is not driven by image quality.

In sum, we found overall excellent reliability of electric field magnitudes, by using a TS design where we acquired two consecutive structural scans in 10 participants on five different 3T Siemens scanners. The magnitudes were similar within individuals in two runs and across scanner sites, with only small differences between target regions. Importantly, variability between runs and sites were not substantially different, indicating that scanner differences were not larger than the measurement bias. Small differences in sequence acquisition procedures did not substantially affect the reliability of electric field magnitudes. Despite our harmonization efforts prior to data acquisition, different scanner software versions and reconstruction algorithms might have contributed to difference in quality metrics which however did not hamper the reliability of simulated tDCS-induced individual electric field magnitudes. In terms of ICC values, our results were consistent with previous studies investigated other imaging sequences, including volumetric measures from structural scans, resting-state imaging, or diffusion tensor imaging [2].

We provide the first TS study delineating the reliability of electric field simulations. Within our ongoing multicenter study, this dataset can be implemented for cross-scanner harmonization of electric field estimates [8] to allow joint data analyses of dose-response relationships. Overall, our evidence of highly reliable electric field estimates provides important ground-work for other multicenter brain stimulation studies as well.

## Authorship statement

The authors confirm that they meet the requirement for authorship.

## CRediT authorship contribution statement

**Dayana Hayek:** Writing – review & editing, Writing – original draft, Visualization, Project administration, Investigation, Formal analysis, Conceptualization. **Daria Antonenko:** Writing – review & editing, Writing – original draft, Visualization, Supervision, Project administration, Investigation, Funding acquisition, Conceptualization. **Ulrike Grittner:** Visualization, Formal analysis. **Axel Thielscher:** Writing – review & editing, Writing – original draft, Funding acquisition, Conceptualization.

