## Supplementary Material for "Multicenter reliability of electric field simulations: Evidence from a traveling-subjects study"

**Supplementary Methods**

*Participants and study design.* The Traveling Subjects study was conducted within the framework of our multicenter study on modulation of brain networks for memory and learning by transcranial electrical brain stimulation (memoslap.de) to evaluate inter-site consistency and data quality across MRI scanners. Ten healthy adults (age range: 22-45 years, M = 33, SD = 8; 5 females) each travelled to five German imaging sites (Berlin, Dortmund, Essen, Greifswald, and Leipzig), where they underwent two consecutive MRI sessions per visit. This traveling-subject design yielded 100 MRI sessions per modality. Participants had no history of neurological or psychiatric disorders. The study was approved by the local ethics committees of all participating institutions, and written informed consent was obtained from all participants prior to data collection.

*MRI data acquisition.* At each site, participants underwent a standardized multimodal MRI protocol including high-resolution T1-weighted and T2-weighted anatomical scans used for personalized head modeling and anatomical reference. All sites used Siemens 3T MRI systems with 64-channel head-neck coils: MAGNETOM Vida (Greifswald, Essen; software version syngo MR XA50), MAGNETOM Prisma or Prisma Fit (Berlin: syngo MR XA30; Dortmund: syngo MR E11), and Prisma_fit (Leipzig: syngo MR E11). Acquisition parameters were harmonized across scanners to ensure comparable image quality. For T1-weighted imaging, we employed a 3D magnetization-prepared rapid gradient-echo (MPRAGE) sequence with the following parameters: 0.9 mm isotropic voxels, TR = 2700 ms, TE = 3.68-3.70 ms, TI = 1090 ms, flip angle = 9°, base resolution = 288, parallel imaging factor (GRAPPA) = 2, and partial Fourier factor = 0.875. For T2-weighted imaging, we used a 3D sampling perfection with application optimized contrasts using different flip angle evolution (SPACE) sequence: 0.9 mm isotropic voxels, TR = 2500 ms, TE = 349 ms, flip angle = 120°, base resolution = 320, number of averages = 2, and 2×2 parallel imaging acceleration. Despite protocol harmonization efforts, systematic differences remained across sites due to different scanner software versions and sequence implementations. Specifically, sites using older software (syngo MR E11: Leipzig, Dortmund) employed different reconstruction algorithms and k-space sampling strategies compared to sites using newer software versions (syngo MR XA30/XA50: Berlin, Greifswald, Essen). For T2 sequences, Leipzig and Dortmund utilized standard GRAPPA parallel imaging with turbo spin-echo variable flip angle reconstruction (225 phase encoding steps), while Berlin, Greifswald, and Essen employed CAIPIRINHA (Controlled Aliasing in Parallel Imaging Results in Higher Acceleration) with optimized 3D encoding (288 phase encoding steps). All structural images underwent standardized quality control using the MRIQC pipeline [1] to assess image quality metrics including entropy focus criterion (EFC), signal-to-noise ratio (SNR), and contrast-to-noise ratio (CNR).

*Head segmentation.* Individual head models were created using the SimNIBS 4.1 “charm” pipeline [2], combining T1- and T2-weighted images for improved tissue contrast. The pipeline produced tetrahedral head meshes including scalp, compact and spongy skull bone, cerebrospinal fluid (CSF), grey and white matter, eyes, and large blood vessels. Each mesh was visually inspected to ensure anatomical accuracy and completeness. We also reconstructed a surface approximating the middle of the cerebellar grey matter by solving the Laplace equation between the grey-white (set to potential 0) and grey-CSF (potential 1) boundaries and extracting the 0.5 isopotential surface.

*ROI definition.* Target regions of interest (ROIs) were defined based on the multicenter study framework described in [3]. Briefly, group-level ROIs for cortical targets (right occipitotemporal cortex, left posterior parietal cortex, left inferior frontal gyrus, left primary motor cortex, right dorsolateral prefrontal cortex, left dorsolateral prefrontal cortex) were defined on fsaverage space (FreeSurfer software) and mapped to individual middle gray matter surfaces using surface-based registration embedded in SimNIBS (Simulation of Non-invasive Brain Stimulation software). The cerebellar ROI (right cerebellum) was defined in MNI space (Montreal Neurological Institute space) and non-linearly transformed to individual space. The resulting subject-specific ROIs served as target regions for electric field extraction.

*Electric field simulation.* Focal tDCS was simulated using SimNIBS v4.1 with a 3×1 center-surround montage at 2 mA, applying the region-specific center-surround radii determined in [3]. Center electrode positions were individualized using the Euclidean optimization algorithm, which minimizes the Euclidean distance between the scalp position and the center of gravity of the target ROI. Electric field magnitudes (V/m) were extracted as the median field strength within each ROI from the individual middle gray matter surface outputs. Standard SimNIBS tissue conductivity values were used throughout.

*Tissue volume calculation.* Volumes of scalp, skull (compact and spongy bone combined), and CSF were extracted both globally and within cylindrical regions of interest (radius 20 mm, depth 40 mm) centered beneath the stimulation sites. Tissue-specific tetrahedra within these cylinders were summed to obtain local volume estimates (in mm³), which were used to assess site- and subject-level anatomical variability influencing current flow.

*Image quality calculation.* Structural image quality was assessed using the MRIQC pipeline [1]. The following metrics were extracted for both T1- and T2-weighted images: entropy focus criterion (EFC), a whole-image metric based on the Shannon entropy of voxel intensities that is sensitive to motion-related artifacts; signal-to-noise ratio (SNR), calculated as the mean signal intensity divided by its standard deviation within gray matter, white matter, and CSF masks; and contrast-to-noise ratio (CNR), defined as the contrast between gray and white matter signal intensities relative to background noise. For a detailed description of metric definitions and computation, see [4].

*Statistical analyses.* Statistical analyses were performed in R (version 4.3.1) using the packages *lme4*, *lmerTest*, *partR2*, and *MuMIn*.

- Variance decomposition of electric field magnitudes. To quantify inter-scanner and scan-rescan reliability of simulated electric field magnitudes, we fitted linear mixed-effects models (LMMs) for each target region (rOTC, lPPC, lIFG, lM1, rCB, rDLPFC, lDLPFC) using *lme4*. The model included random intercepts for subject, site, and run nested within site, decomposing total variance into between-subject, between-site, within-site (scan-rescan), and residual components. Intraclass correlation coefficients (ICCs) reflecting inter-scanner and scan-rescan reliability were derived from these variance components and accompanied by 95% confidence intervals obtained via parametric bootstrapping (1,000 resamples). To test whether between-site variance differed significantly from within-site variance, we generated bootstrap distributions of the difference between the corresponding ICC estimates, 95%CI and derived two-sided p-values.
- Within-person variance decomposition. To examine the contribution of intra-individual tissue segmentation variability to electric field magnitude variability, all variables were decomposed into within-person components by subtracting each individual's mean, computed across all sessions and sites, from their observation-level values. This person-mean centering isolates variability attributable to measurement occasion rather than stable individual differences. We divided all tissue volumes by 10,000 before running the analysis to bring all the numbers into a similar range. For tissue volumes (GM, CSF, bone, scalp), the sign of the deviation was inverted for scalp, bone, and CSF, such that higher deviation values consistently reflect conditions expected to increase field magnitude (i.e., less tissue between electrode and cortex).
- For each target region separately, we fitted LMMs predicting within-person magnitude deviations: a model including within-person deviations of all four tissue types as fixed effects (*magnitude_within* ~ *bone_within* + *scalp_within* + *csf_within* + *gm_within* + (1|subject) + (1|site)). The joint contribution of the four tissue volume predictors to the marginal variance in magnitude was quantified as semi-partial R² and 95%CI using the *partR2*.
- Image quality as a predictor of segmentation variability. To assess whether intra-individual variability in structural image quality explained segmentation variability, we fitted analogous LMMs for each tissue type (bone, scalp, CSF, GM) as outcome, with within-person deviations of T2 EFC and T1 EFC as fixed-effect predictors: *tissue_within* ~ *t2_within* + *t1_within* + (1|subject) + (1|site). The joint semi-partial R² of T2 and T1 EFC deviations was estimated via *partR2*.

**Supplementary Results**

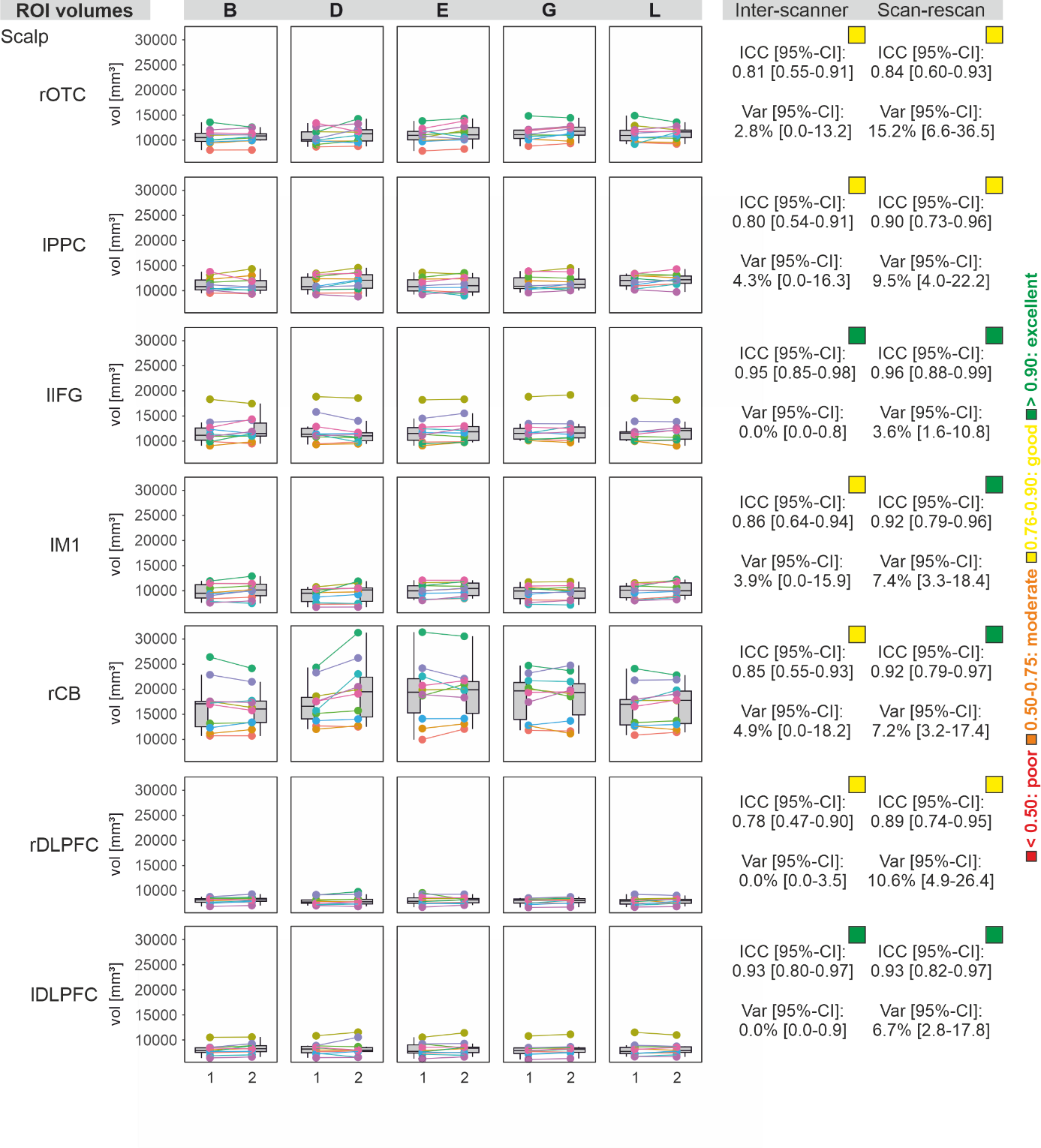

**Supplementary Figure S1.** Scalp volumes, extracted above the regions-of-interest (selected based on our multicenter study memoslap.de). The figures show two consecutive scans/runs for each of the 10 subjects (each in a different color) at five scanner sites (B, Berlin; D, Dortmund; E, Essen; G, Greifswald; L, Leipzig). Right columns show intra-class correlation coefficients (ICC) and variance components (Var) between sites (inter-scanner) and runs (scan-rescan). Colored squares above the ICC values additionally illustrate whether reliability was poor (ICC < 0.50, red), moderate (0.50-0.75, orange), good (0.76-0.90, yellow), or excellent (> 0.90, green). rOTC, right occipitotemporal cortex. lPPC, left posterior parietal cortex. lIFG, left inferior frontal cortex. lM1, left motor cortex. rCB, right cerebellum. r/lDLPFC, right/left dorsolateral prefrontal cortex.

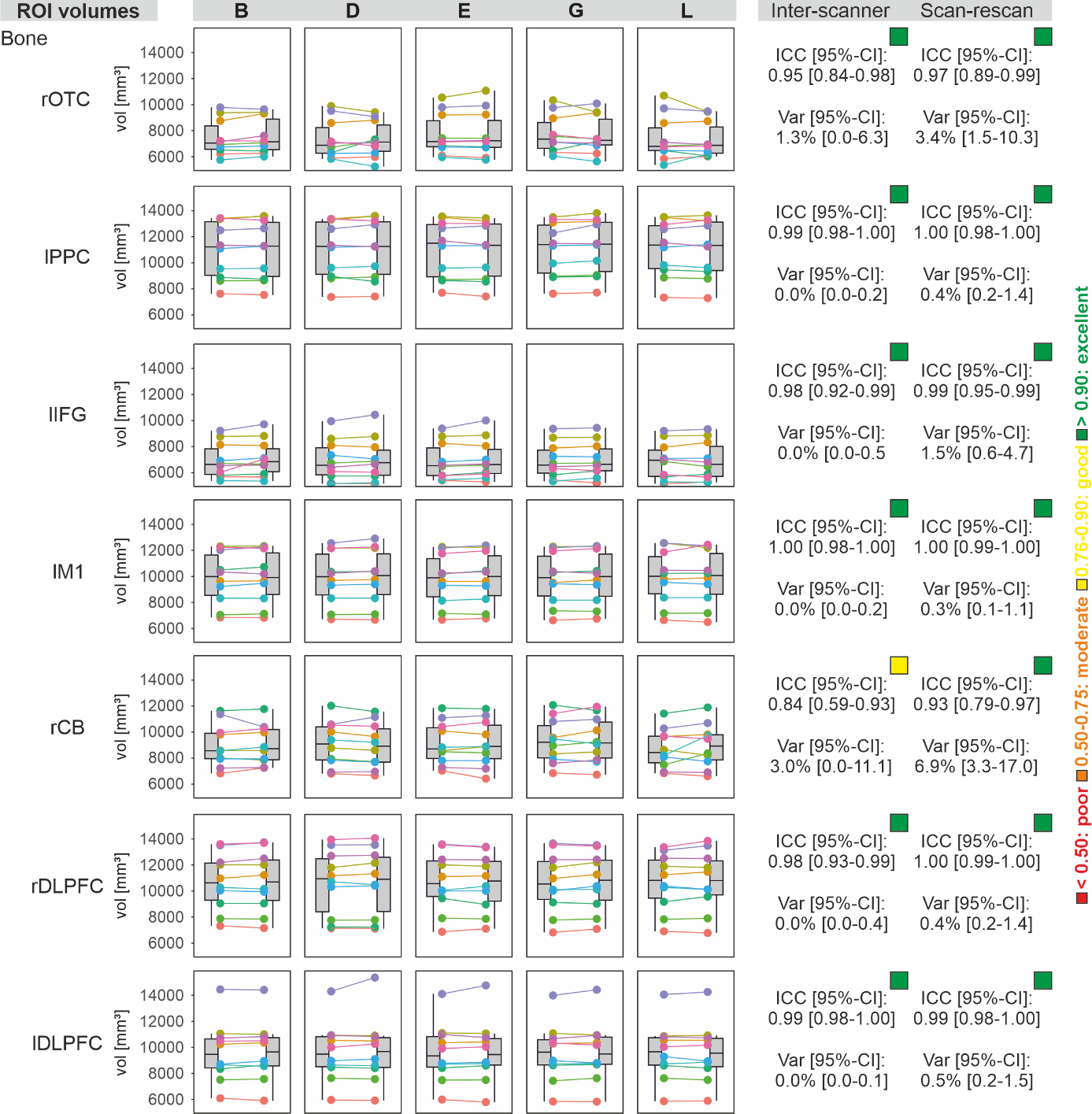

**Supplementary Figure S2.** Bone volumes, extracted above the regions-of-interest (selected based on our multicenter study memoslap.de). The figures show two consecutive scans/runs for each of the 10 subjects (each in a different color) at five scanner sites (B, Berlin; D, Dortmund; E, Essen; G, Greifswald; L, Leipzig). Right columns show intra-class correlation coefficients (ICC) and variance components (Var) between sites (inter-scanner) and runs (scan-rescan). Colored squares above the ICC values additionally illustrate whether reliability was poor (ICC < 0.50, red), moderate (0.50-0.75, orange), good (0.76-0.90, yellow), or excellent (> 0.90, green). rOTC, right occipitotemporal cortex. lPPC, left posterior parietal cortex. lIFG, left inferior frontal cortex. lM1, left motor cortex. rCB, right cerebellum. r/lDLPFC, right/left dorsolateral prefrontal cortex.

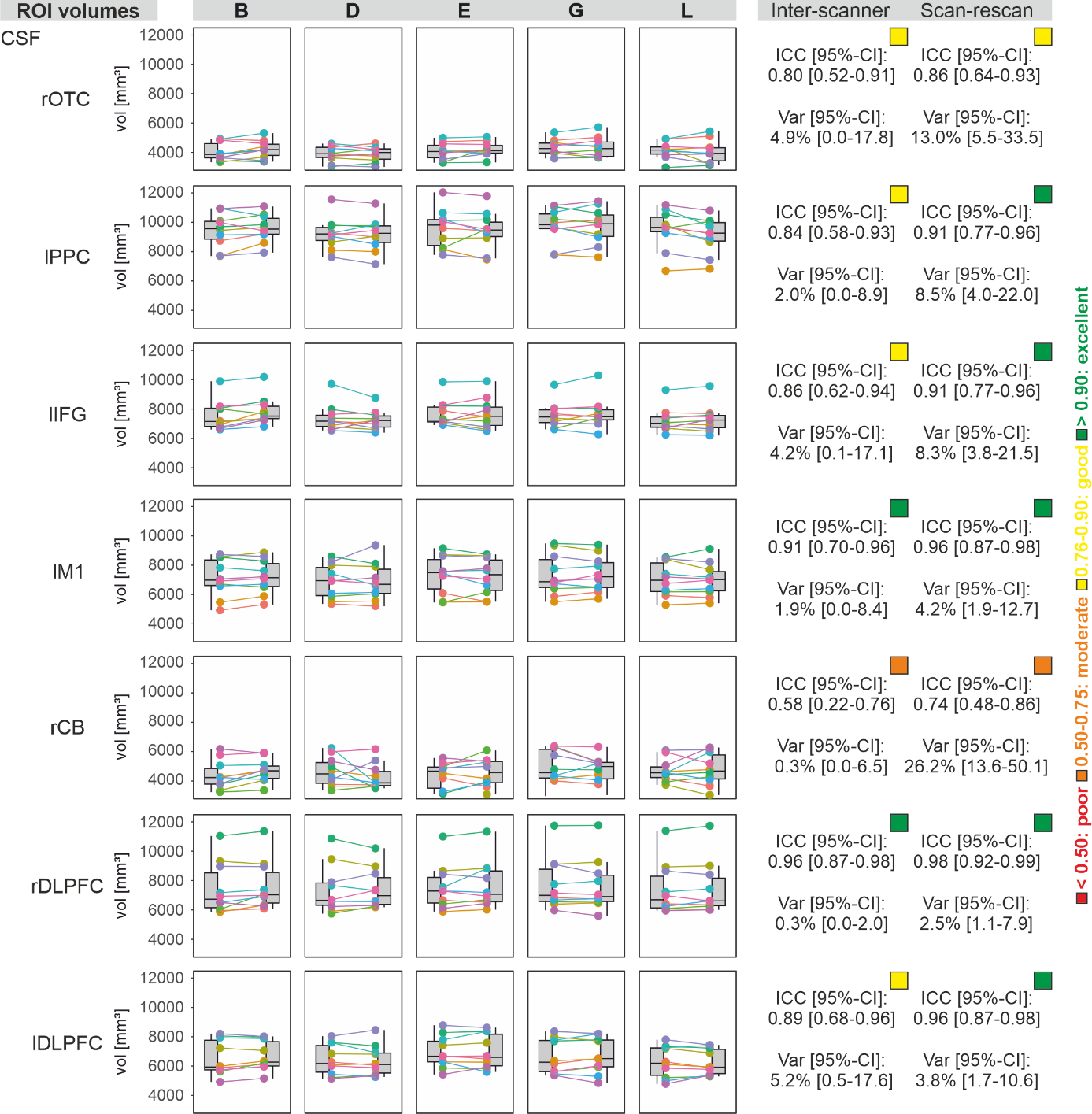

**Supplementary Figure S3.** CSF volumes, extracted above the regions-of-interest (selected based on our multicenter study memoslap.de). The figures show two consecutive scans/runs for each of the 10 subjects (each in a different color) at five scanner sites (B, Berlin; D, Dortmund; E, Essen; G, Greifswald; L, Leipzig). Right columns show intra-class correlation coefficients (ICC) and variance components (Var) between sites (inter-scanner) and runs (scan-rescan). Colored squares above the ICC values additionally illustrate whether reliability was poor (ICC < 0.50, red), moderate (0.50-0.75, orange), good (0.76-0.90, yellow), or excellent (> 0.90, green). rOTC, right occipitotemporal cortex. lPPC, left posterior parietal cortex. lIFG, left inferior frontal cortex. lM1, left motor cortex. rCB, right cerebellum. r/lDLPFC, right/left dorsolateral prefrontal cortex.

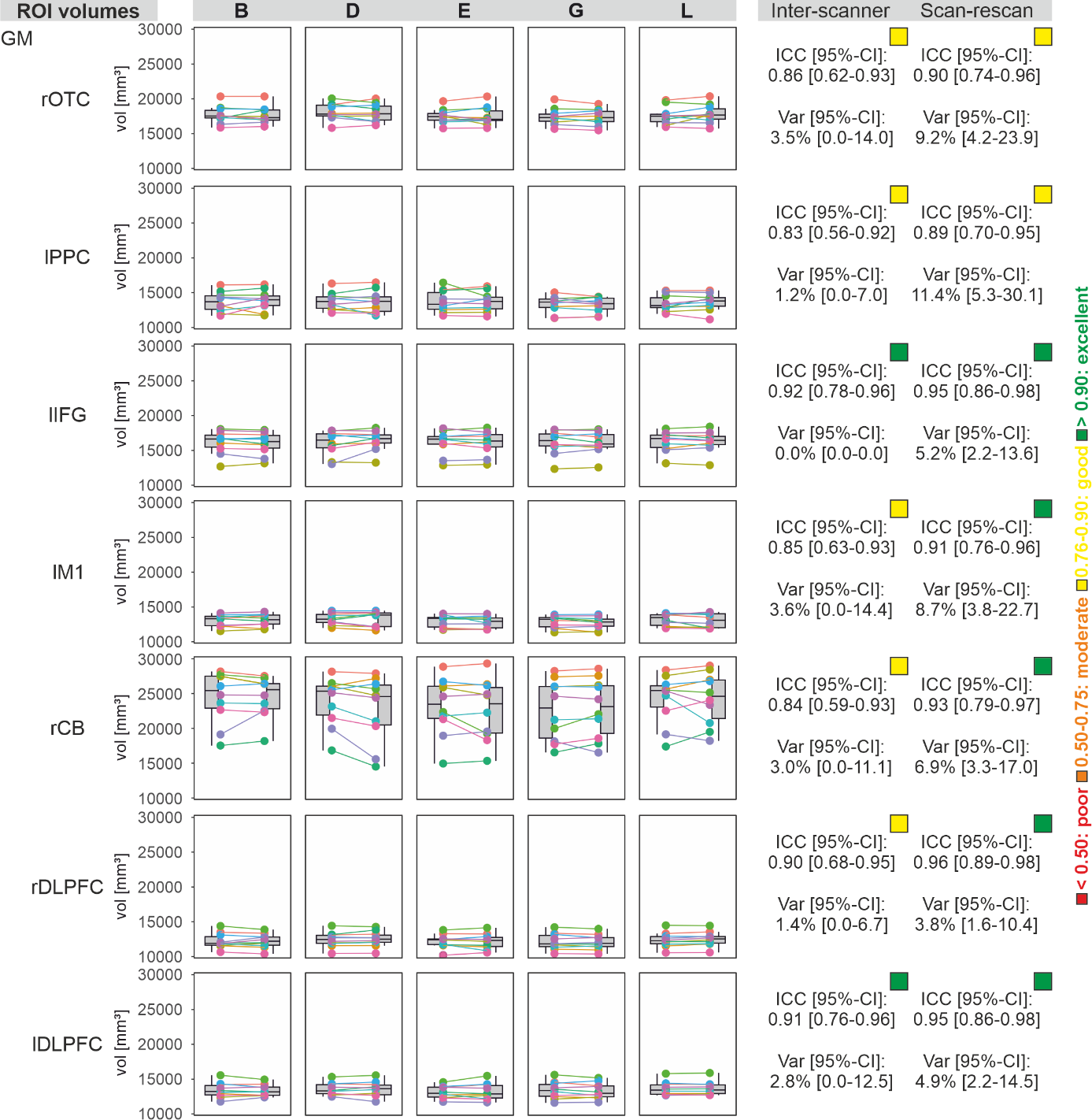

**Supplementary Figure S4.** GM volumes, extracted above the regions-of-interest (selected based on our multicenter study memoslap.de). The figures show two consecutive scans/runs for each of the 10 subjects (each in a different color) at five scanner sites (B, Berlin; D, Dortmund; E, Essen; G, Greifswald; L, Leipzig). Right columns show intra-class correlation coefficients (ICC) and variance components (Var) between sites (inter-scanner) and runs (scan-rescan). Colored squares above the ICC values additionally illustrate whether reliability was poor (ICC < 0.50, red), moderate (0.50-0.75, orange), good (0.76-0.90, yellow), or excellent (> 0.90, green). rOTC, right occipitotemporal cortex. lPPC, left posterior parietal cortex. lIFG, left inferior frontal cortex. lM1, left motor cortex. rCB, right cerebellum. r/lDLPFC, right/left dorsolateral prefrontal cortex.

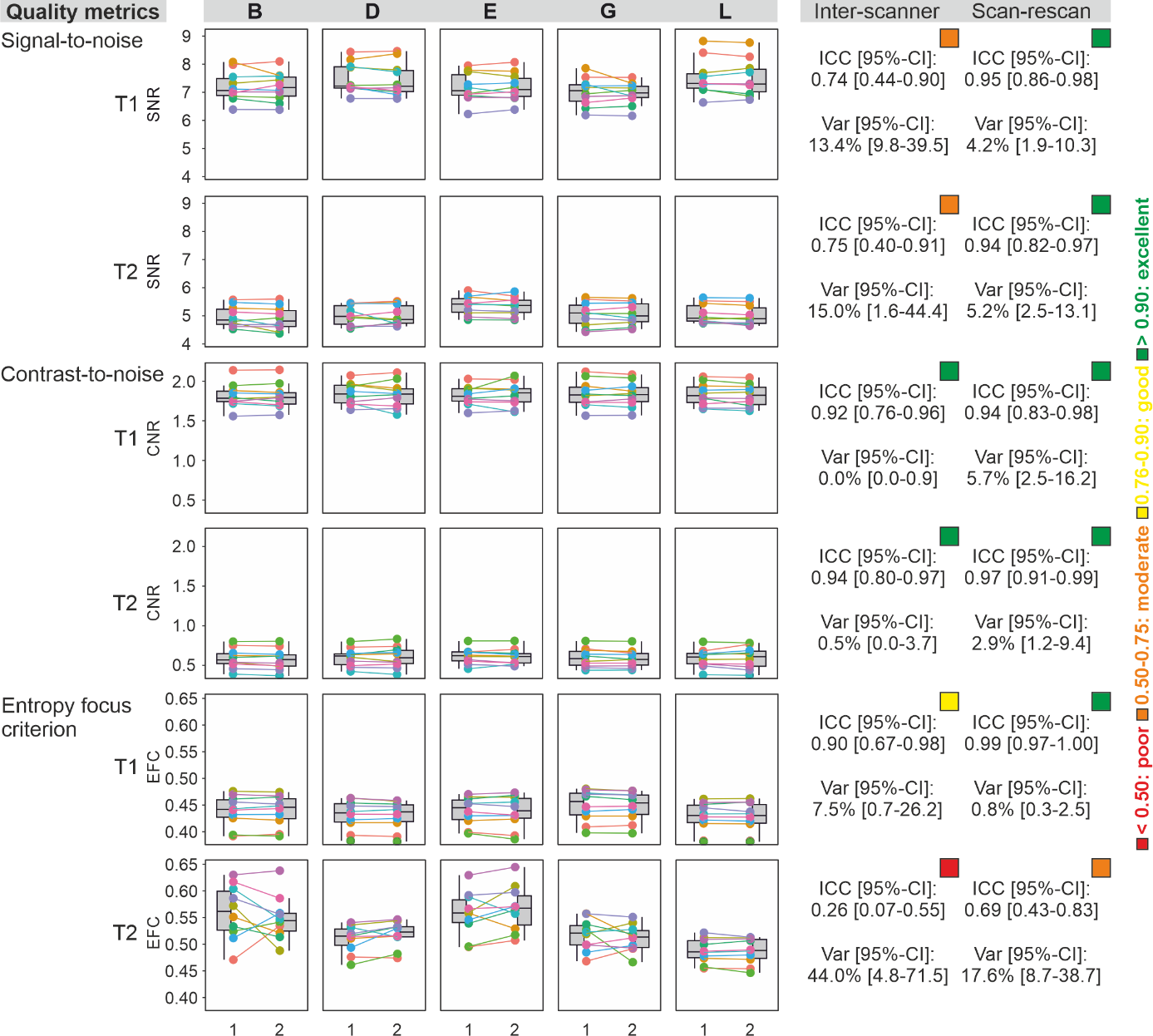

**Supplementary Figure S5.** Quality metrics for structural images. We extracted signal-to-noise ratios (SNR), contrast-to-noise ratio (CNR), and entropy focus criterion (EFC) for T1- and T2-weighted images. The figures show two consecutive scans/runs for each of the 10 subjects (each in a different color) at five scanner sites (B, Berlin; D, Dortmund; E, Essen; G, Greifswald; L, Leipzig). Right columns show intra-class correlation coefficients (ICC) and variance components (Var) between sites (inter-scanner) and runs (scan-rescan). Colored squares above the ICC values additionally illustrate whether reliability was poor (ICC < 0.50, red), moderate (0.50-0.75, orange), good (0.76-0.90, yellow), or excellent (> 0.90, green).
