## Supplementary Table S1 for "Multicenter reliability of electric field simulations: Evidence from a traveling-subjects study"

| **Marginal R² decomposition — volume predictors (within-person deviations) of magnitude (within-person deviations),**  Formula: magnitude_within ~ bone_within + scalp_within + csf_within + gm_within + (1 \| subject) + (1\|site) | | | | |
| --- | --- | --- | --- | --- |
| **Modality** | **Part. R², 95%CI (variance in within-person magnitude deviation explained by within-person volume deviations)** | **variables** | **β, 95% CI** | **p** |
| **right occipitotemporal cortex (rOTC)** | | | | |
|  | 0.44 [0.35, 0.63] | Bone (within) | -0.08 [-0.19, 0.02] | 0.117 |
|  |  | Scalp (within) | 0.18 [0.13, 0.23] | <0.001 |
|  |  | CSF (within) | 0.19 [0.06, 0.31] | 0.003 |
|  |  | GM (within) | 0.04 [0.12, 0.30] | 0.296 |
| **left posterior parietal cortex (lPPC)** | | | | |
|  | 0.46 [0.35, 0.61] | Bone (within) | 0.06 [-0.03, 0.14] | 0.206 |
|  |  | Scalp (within) | 0.10 [0.07, 0.13] | <0.001 |
|  |  | CSF (within) | 0.05 [0.01, 0.09] | 0.016 |
|  |  | GM (within) | -0.02 [-0.06, 0.02] | 0.373 |
| **left inferior frontal gyrus (lIFG)** | | | | |
|  | 0.52 [0.41, 0.64] | Bone (within) | 0.01 [-0.05, 0.07] | 0.800 |
|  |  | Scalp (within) | 0.03 [0.00, 0.06] | 0.028 |
|  |  | CSF (within) | 0.14 [0.10, 0.18] | <0.001 |
|  |  | GM (within) | 0.03 [-0.01, 0.07] | 0.096 |
| **left primary motor cortex (lM1)** | | | | |
|  | 0.59 [0.50, 0.72] | Bone (within) | -0.06 [-0.16, 0.04] | 0.220 |
|  |  | Scalp (within) | 0.12 [0.10, 0.15] | <0.001 |
|  |  | CSF (within) | 0.13 [0.09, 0.17] | <0.001 |
|  |  | GM (within) | -0.01 [-0.06, 0.03] | 0.542 |
| **right cerebellum (rCB)** | | | | |
|  | 0.71 [0.65, 0.80] | Bone (within) | 0.07 [0.00, 0.14] | 0.037 |
|  |  | Scalp (within) | 0.11 [0.09, 0.14] | <0.001 |
|  |  | CSF (within) | 0.22 [0.17, 0.26] | <0.001 |
|  |  | GM (within) | -0.08 [-0.11, -0.05] | <0.001 |
| **right dorsolateral prefrontal cortex (rDLPFC)** | | | | |
|  | 0.57 [0.49, 0.68] | Bone (within) | 0.04 [-0.01, 0.08] | 0.107 |
|  |  | Scalp (within) | 0.16 [0.13, 0.20] | <0.001 |
|  |  | CSF (within) | 0.12 [0.08, 0.16] | <0.001 |
|  |  | GM (within) | -0.01 [-0.06, 0.04] | 0.623 |
| **left dorsolateral prefrontal cortex (lDLPFC)** | | | | |
|  | 0.59 [0.49, 0.69] | Bone (within) | -0.03 [-0.11, 0.05] | 0.449 |
|  |  | Scalp (within) | 0.11 [0.07, 0.15] | <0.001 |
|  |  | CSF (within) | 0.09 [0.04, 0.15] | 0.002 |
|  |  | GM (within) | 0.09 [0.03, 0.16] | 0.005 |
| Part. R² = semi-partial R² (marginal) for the joint contribution of all within-person volume predictors, estimated via partR2 (95% CI).  R²m = total marginal R²; R²c = total conditional R². LRT = likelihood ratio test vs. null model (m0). rOTC = right occipitotemporal cortex;  lPPC = left posterior parietal cortex; lIFG = left inferior frontal gyrus; lM1 = left primary motor cortex; rCB = right cerebellum;  rDLPFC = right dorsolateral prefrontal cortex; lDLPFC = left dorsolateral prefrontal cortex. | | | | |
