## Supplementary Table S2 for "Multicenter reliability of electric field simulations: Evidence from a traveling-subjects study"

**Supplementary Table S3**

| **Marginal R² decomposition — quality within-person deviation predictors of volume within-person deviation outcomes**  Outcomes: bone/scalp/csf/gm (within) ~ t2_within + t1_within + (1\|subject) + (1\|site) | |
| --- | --- |
| **Formula** | **Part. R², , 95%CI (variance in within-person volume deviation explained by within-person quality deviations)** |
| **right occipitotemporal cortex (rOTC)** | |
| bone_within ~ t2_within + t1_within + (1 \| subject) + (1 \| site) | 0.07 [0.02, 0.25] |
| scalp_within ~ t2_within + t1_within + (1 \| subject) + (1 \| site) | 0.01 [0.00, 0.20] |
| csf_within ~ t2_within + t1_within + (1 \| subject) + (1 \| site) | 0.04 [0.00, 0.20] |
| gm_within ~ t2_within + t1_within + (1 \| subject) + (1 \| site) | 0.10 [0.01, 0.34] |
| **left posterior parietal cortex (lPPC)** | |
| bone_within ~ t2_within + t1_within + (1 \| subject) + (1 \| site) | 0.06 [0.01, 0.15] |
| scalp_within ~ t2_within + t1_within + (1 \| subject) + (1 \| site) | 0.15 [0.06, 0.29] |
| csf_within ~ t2_within + t1_within + (1 \| subject) + (1 \| site) | 0.16 [0.08, 0.30] |
| gm_within ~ t2_within + t1_within + (1 \| subject) + (1 \| site) | 0.01 [0.00, 0.14] |
| **left inferior frontal gyrus (lIFG)** | |
| bone_within ~ t2_within + t1_within + (1 \| subject) + (1 \| site) | 0.01 [0.00, 0.06] |
| scalp_within ~ t2_within + t1_within + (1 \| subject) + (1 \| site) | 0.01 [0.00, 0.12] |
| csf_within ~ t2_within + t1_within + (1 \| subject) + (1 \| site) | 0.06 [0.01, 0.32] |
| gm_within ~ t2_within + t1_within + (1 \| subject) + (1 \| site) | 0.03 [0.01, 0.15] |
| **left primary motor cortex (lM1)** | |
| bone_within ~ t2_within + t1_within + (1 \| subject) + (1 \| site) | 0.07 [0.01, 0.22] |
| scalp_within ~ t2_within + t1_within + (1 \| subject) + (1 \| site) | 0.00 [0.00, 0.18] |
| csf_within ~ t2_within + t1_within + (1 \| subject) + (1 \| site) | 0.16 [0.05, 0.31] |
| gm_within ~ t2_within + t1_within + (1 \| subject) + (1 \| site) | 0.18 [0.08, 0.34] |
| **right cerebellum (rCB)** | |
| bone_within ~ t2_within + t1_within + (1 \| subject) + (1 \| site) | 0.12 [0.05, 0.25] |
| scalp_within ~ t2_within + t1_within + (1 \| subject) + (1 \| site) | 0.02 [0.01, 0.23] |
| csf_within ~ t2_within + t1_within + (1 \| subject) + (1 \| site) | 0.02 [0.00, 0.12] |
| gm_within ~ t2_within + t1_within + (1 \| subject) + (1 \| site) | 0.00 [0.00, 0.16] |
| **right dorsolateral prefrontal cortex (rDLPFC)** | |
| bone_within ~ t2_within + t1_within + (1 \| subject) + (1 \| site) | 0.01 [0.00, 0.09] |
| scalp_within ~ t2_within + t1_within + (1 \| subject) + (1 \| site) | 0.04 [0.01, 0.17] |
| csf_within ~ t2_within + t1_within + (1 \| subject) + (1 \| site) | 0.17 [0.07, 0.34] |
| gm_within ~ t2_within + t1_within + (1 \| subject) + (1 \| site) | 0.24 [0.11, 0.38] |
| **left dorsolateral prefrontal cortex (lDLPFC)** | |
| bone_within ~ t2_within + t1_within + (1 \| subject) + (1 \| site) | 0.01 [0.00, 0.09] |
| scalp_within ~ t2_within + t1_within + (1 \| subject) + (1 \| site) | 0.03 [0.00, 0.15] |
| csf_within ~ t2_within + t1_within + (1 \| subject) + (1 \| site) | 0.08 [0.01, 0.29] |
| gm_within ~ t2_within + t1_within + (1 \| subject) + (1 \| site) | 0.16 [0.02, 0.37] |
| Part. R² = semi-partial R² (marginal) for the joint contribution of t2_within and t1_within, estimated via partR2 (100 bootstrap resamples, 95% CI). R²m = total marginal R²; R²c = total conditional R². LRT = likelihood ratio test vs. null model (m0). rOTC = right occipitotemporal cortex; lPPC = left posterior parietal cortex; lIFG = left inferior frontal gyrus; lM1 = left primary motor cortex; rCB = right cerebellum; rDLPFC = right dorsolateral prefrontal cortex; lDLPFC = left dorsolateral prefrontal cortex. | |
